# Exploring the maturation of a monocytic cell line using self-organizing maps of single-cell Raman spectra

**DOI:** 10.1101/2020.05.30.124552

**Authors:** Sayani Majumdar, Mary L Kraft

## Abstract

Phorbol myristate acetate (PMA)-differentiated THP-1 cells are routinely used in lieu of primary macrophages to study macrophage polarization during host-pathogen interactions and disease progression. The phenotypes of the THP-1 macrophages are influenced by the level and duration of PMA stimulation, and possibly also by the presence of adhesion factors. Here, we use self-organizing maps (SOMs) of single-cell Raman spectra to probe the effects of PMA stimulation conditions and adhesion factors on THP-1 cell differentiation. Raman spectra encoding for biochemical composition were acquired from individual cells on substrates coated with fibronectin or poly-L-lysine before and after stimulation with 20 nM or 200 nM PMA for two different time intervals. SOMs that show the extent of spectral dissimilarity were constructed. For all stimulation conditions, the SOMs indicated the spectra acquired from cells after 3 d treatment had diverged from those of untreated cells. The SOMs also showed treatment with 200 nM PMA for 3 d produced both fully and partially differentiated cells on both adhesion factors, whereas the outcome of 20 nM PMA treatment for 3 d depended on the adhesion factor. Treatment of THP-1 cells on fibronectin with 20 nM PMA for 3 d produced both partially and fully differentiated cells, but this treatment induced an intermediate stage of differentiation when applied to THP-1 cells on poly-L-lysine. Thus, the transition of THP-1 monocytes into macrophage-like cells may be modulated by integrin-binding interactions. Furthermore, the composite of culture and stimulation conditions may confound the comparison of results from separate studies.

## INTRODUCTION

Macrophages comprise a phenotypically diverse population of cells with pivotal roles in host defense, tissue remodeling and homeostasis.^1^ In their naïve or resting state, tissue-resident macrophages are responsible for the clearance of cellular debris resulting from tissue reorganization and apoptosis. Exogenous stimuli can transiently reprogram macrophages towards the pro-inflammatory classically activated phenotype or the anti-inflammatory alternatively activated phenotype, also called wound-healing macrophages, each with their own distinct functions. These activated macrophages have been implicated in diseases caused by persistent inflammation and tissue remodeling, such as atherosclerosis,^2^ rheumatoid arthritis,^3^ and tissue fibrosis.^4^ Regulatory macrophages, which may be a sub-type of alternatively activated macrophages, have also emerged as an increasingly important player in the progression of tumors,^5^ Studies of macrophage behavior under various pathophysiological conditions are crucial to designing effective treatments. Macrophages obtained by the differentiation of primary human peripheral blood mononuclear cells (PBMC) derived-monocytes may be used in these studies. However, the variability in genotype, limited availability, and ethical issues surrounding use and handling of PBMC derived monocytes and macrophages have led to the extensive use of monocytic cell lines, especially the THP-1 cell line, in studies of monocyte and macrophage function and activities.^6^

The treatment of THP-1 cells with phorbol 12-myristate 13-acetate (PMA) induces their differentiation into macrophage-like cells,^7, 8^ referred to as THP-1 macrophages, that exhibit many hallmarks of resting macrophages *in vivo*. This includes stronger adherence to the substrate, higher phagocytic capacity, larger numbers of ribosomes, lysosomes and mitochondria, and the ability to polarize into different types of activated macrophage cells following exposure to appropriate stimuli.^5, 9, 10^ The extent of THP-1 monocyte differentiation to the resting macrophage phenotype is influenced by the PMA concentration used for stimulation,^5, 8, 9, 11–14^ and whether the cells are allowed to rest in PMA-free medium for at least 24 h following PMA treatment.^6^ Furthermore, monocytes interact extensively with extracellular matrix (ECM) proteins before and during their differentiation in various physiological and pathological contexts in vivo. Additionally, the differentiation of human PBMC derived monocytes to macrophages is strongly accelerated by fibronectin.^15^ Therefore, fibronectin may also modulate the PMA-induced differentiation of THP-1 cells. Such potential differences in the degree of differentiation are reflected by differences in receptor expression, functional characteristics, and other macrophage markers, which could complicate efforts to reconcile the findings of different studies.

In this study, we investigate the combined effects of PMA stimulation conditions and fibronectin on the extent of THP-1 monocyte differentiation to a macrophage-like phenotype at specified time points. For comparison with fibronectin, poly-L-lysine was selected because it promotes nonspecific cell adhesion to the substrate via electrostatic interactions, whereas the effects of fibronectin can be attributed to specific receptor-fibronectin binding interactions. We examined the effects of fibronectin and poly-L-lysine on the dynamics of THP-1 cell differentiation induced by two PMA concentrations that differ by an order of magnitude (20 and 200 nM) but still lie in the range of concentrations that have been used to differentiate THP-1 cells (8.1 to 648.5 nM or 5 to 400 ng/ml).^5, 8, 9, 11–14^ We analyzed the THP-1 cells immediately after PMA treatment and after resting the PMA-treated cells in PMA-free medium for five days because the inclusion of this rest period is reported to increase THP-1 cell differentiation.^8^

To assess the extent of THP-1 cell differentiation to the macrophage-like phenotype, we used self-organizing maps (SOMs) of single-cell Raman spectra. Because Raman microspectroscopy is a non-invasive, label-free approach, it enables acquiring spectra from individual cells at several time-points over the course of treatment. The individual peaks in a single-cell spectrum are associated with specific chemical bonds and functional groups, and the compilation of these peaks encodes for the cell’s biochemical composition. Because cellular composition is intimately linked to function, the cell’s phenotype is encrypted in its spectral fingerprint.^16, 17^ To visualize the extent of the spectral, and thus, phenotypic differences between individual cells subjected to different PMA stimulation regimens and adhesion factors, we constructed SOMs of the spectra^18^. These SOMs depict the amounts of spectral variation between every cell in the experiment as the distances between their positions on the map^19^ while preserving the interrelationships between all the cell spectra.^20^

## EXPERIMENTAL

### Culture and differentiation of THP-1 cells

Human monocytic THP-1 cells (ATCC) were cultured in suspension in RPMI-1640 medium (Gibco) supplemented with 10% heat-inactivated FBS (Gibco or Sigma) and 1% antibiotic/antimycotic (10000 U/ml penicillin, 10 mg/ml streptomycin, 25 μg/ml amphotericin B, Sigma). For compatibility with Raman spectroscopy, glass substrates coated with a silicon dioxide-protected gold mirror were used as cell supports. Prior to seeding cells, the substrates were passively coated with fibronectin (Corning) or poly-L-lysine (Sigma-Aldrich). Briefly, substrates were incubated with 150 μl of a fibronectin (50 μg/ml) for 1 h or poly-L-lysine (1 mg/ml in water) for 30 min, washed with PBS and dried for 15 - 20 min in a laminar flow hood. Cells were plated on these coated substrates at densities of 3 - 5 × 10^5^ cells/ml. Differentiation into a macrophage-like phenotype was induced by treatment with phorbol 12-myristate 13-acetate (PMA, Sigma-Aldrich,) following the 8-day protocol reported by Daigneault et al.^8^ Cells were incubated with medium containing 20 nM or 200 nM PMA for 72 h, after which the cells were rinsed with PBS to remove PMA and cultured in PMA-free RPMI medium for 5 more days. Most cells were adherent after the 72 h PMA treatment.

### Raman spectroscopy of individual cells

For spectral acquisition, substrates with cells were transferred to a 60-mm-diameter tissue culture dish (Falcon) containing phenol red-free RPMI-1640 medium supplemented with serum and HEPES buffer (Gibco) for better pH control under ambient conditions. Samples were maintained at 37 °C with a heating stage constructed following Heidemann et al.^21^ Raman spectra were acquired of individual THP-1 cells with a confocal Raman imaging system (Horiba LabRAM HR 3D) equipped with a 330 mW 785 nm diode-based laser source (Crystalaser Reno). An Olympus 60X, NA 1.0 water-dipping objective with a working distance of 2 mm was used to focus on a spot of approximately 10 – 15-μm-diameter on each cell. The laser intensity was set to 25% of the full intensity, the pinhole size to 500 mm, the slit size to 100 mm, and the grating to 300 grooves per mm for all measurements. For each cell, a Raman spectrum was recorded in the fingerprint region (600 - 1750 cm^−1^) with a spectral integration time of 30 s. Approximately 35-40 spectra were acquired from THP-1 cells on day 0 (untreated), day 3 (3 d post-PMA) and day 8 (8 d post-PMA) from each fibronectin or poly-L-lysine-coated substrate.

### Preprocessing of Raman spectra

Raman spectra were individually inspected for cosmic spikes and irregular peaks, which were removed using a cosmic spike-removal function within LabSpec 5. Spectra were manually aligned to the phenylalanine peak at 1004 cm^−1^. The fluorescence baseline and the background due to silicon dioxide, fibronectin, or poly-L-lysine was reduced by using the algorithm described by Beier et al.^22^ Using the PLS toolbox, offset alignment (slack = 5) to a randomly chosen spectrum was performed to correct any residual peak misalignment. The Savitzky-Golay algorithm was applied (2^nd^ derivative, 2^nd^ degree polynomial, 25 point-window) to each spectrum to remove the baseline and reduce noise through smoothing. Spectra were normalized so the area under each spectrum equaled one, and subsequently mean-centered.

### Self-organizing map construction

Spectral preprocessing was performed using LabSpec 5 (Horiba Scientific) and the PLS Toolbox (v.8.11.0, Eigenvector Research, Inc.) run in MATLAB (v.8.3.0.532, R2014a, MathWorks, Inc.). The preprocessed spectra were used to train SOMs with the SOM Toolbox (v.2.0 www.cis.hut.fi/projects/somtoolbox/download) run in MATLAB (v.8.3.0.532, 2014a, MathWorks, Inc.). All maps were constructed using the default settings: linear initialization, hexagonal topology, and map dimensions determined according to the two largest eigenvalues of the covariance matrix of the spectral data.^23^ SOM quality was assessed by computing the mean quantization error (mqe) and topographical error (te) (Table S1). The mqe measures how well each cell spectrum matches its BMU, and it is calculated as the root mean square of the differences in peak intensities between the two spectra. The te represents the proportion of the cell spectra for which the next best-matching map unit is not adjacent to the BMU. Maps with low te better represent the similarities between spectra as their relative positions on the map than those with higher te. For visualization, hit histograms and U-matrices were generated. To facilitate locating specific map units in the hit histogram, units without hits are labeled with the number and letter denoting their row and column, respectively. The magnitude of dissimilarity between the map units in the hit histogram are encrypted in the U-matrix. The color-coded hexagons in the U-matrix represent either the average degree of dissimilarity between one map unit in the hit histogram and all of its neighboring map units (labeled with a letter), or the degree of dissimilarity between two adjacent map units (marked with a vertical or angled line).

Component planes, which show the Raman signal intensities of each wave number in the weight vector of every map unit in the SOM, were also generated for four peaks related to lipids, proteins and genetic material. The peaks are: 784 cm^−1^ for DNA and RNA (viz. O-P-O stretching in DNA backbone, U, T, C ring breathing); 811 cm^−1^ for RNA (O-P-O stretching in RNA backbone); 1449 cm^−1^ for mainly lipids and proteins (C-H deformation); 1655 cm^−1^ for proteins and lipids (C=O stretching from amide I in α-helix, C=C stretching).^24^

## RESULTS

Raman spectra were acquired from THP-1 cells on substrates coated with each adhesion factor before PMA treatment, immediately after exposure to PMA for 3 d, and after culturing the PMA-treated cells in PMA-free medium for an additional 5 d to ensure complete differentiation (Figure 1a, c).^8^ Treatment with PMA produced distinct morphological changes in the cells, including adherence to the substrate and an overall increase in apparent size and cell spreading (Figure 1b).

**Figure 1:**
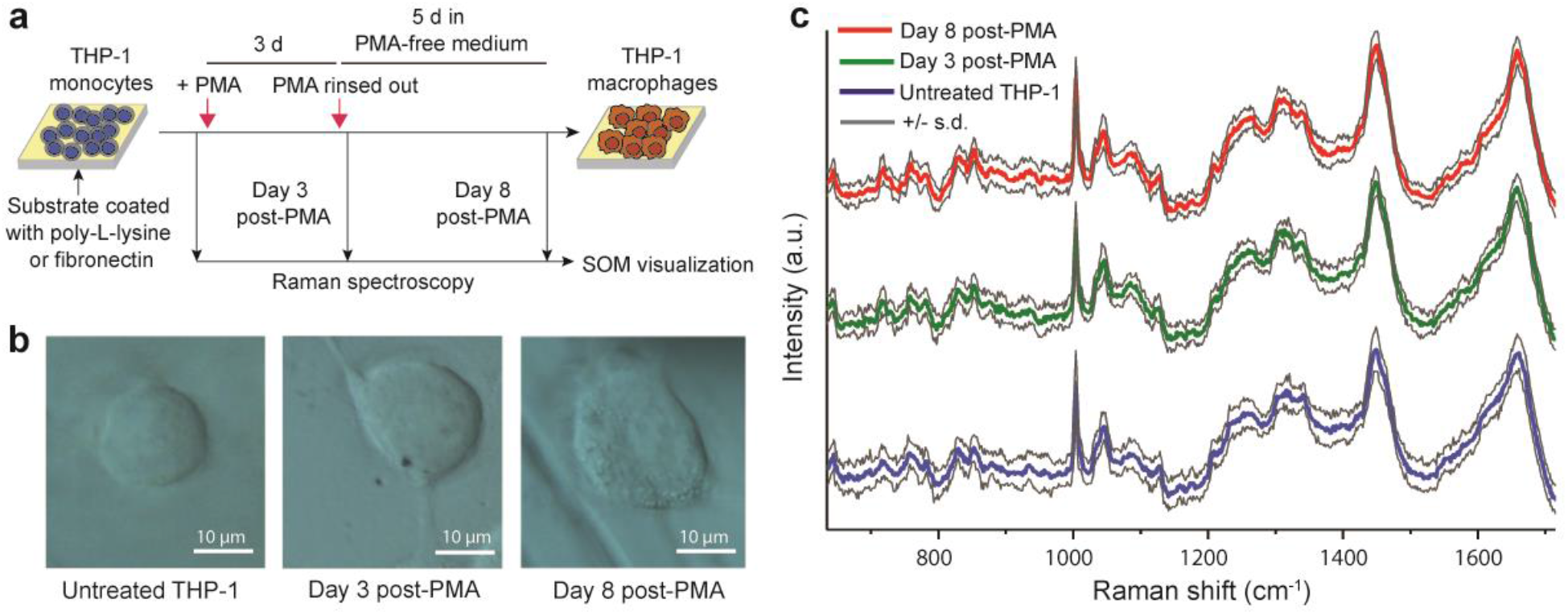
(a) Scheme shows the protocol used for inducing differentiation in THP-1 cells and the time points when Raman microspectroscopy was performed. (b) Morphology of THP-1 cells before and after treatment with PMA. (c) Average (n = 30-40) baseline-corrected Raman spectra of untreated THP-1 cells (blue), 3 d post-PMA cells (green) and 8 d post-PMA cells (red) on poly-L-lysine-coated substrates. Grey lines show 1 standard deviation (s.d.). Spectra were aligned to the phenylalanine peak at 1004 cm^−1^, processed with a weighted least-squares method to remove the baseline, normalized, and offset for clarity.

### Resting THP-1 cells after PMA treatment increases differentiation on poly-L-lysine

An SOM was constructed using the Raman spectra acquired from THP-1 cells on a poly-L-lysine-coated substrate before PMA treatment (untreated THP-1), after treatment with 200 nM PMA for 3 d (3 d post-PMA), or after treatment with 200 nM PMA for 3 d followed by 5 days of rest in PMA-free medium (8 d post-PMA). The resulting SOM (Figure 2a - d) consists of hexagons, or map units, each of which represents a different Raman spectrum. The spectra represented by neighboring map units are more similar to each other than the spectra of units that are further apart on the map. Each cell spectrum is assigned to its best matching unit (BMU), which is the map unit that represents the Raman spectrum most similar to the cell spectrum. The hit histograms (Figure 2a – d) encode the number of times that each map unit is the BMU for an untreated THP-1 cell spectrum, a 3 d post-PMA cell spectrum, or an 8 d post-PMA cell spectrum in the size of the blue, green, and red hexagons, respectively, that are overlaid on the map units. The amount of dissimilarity between adjacent map units in the hit histogram are encoded by the color of the corresponding hexagon in the U-matrix (Figure 2e).

**Figure 2:**
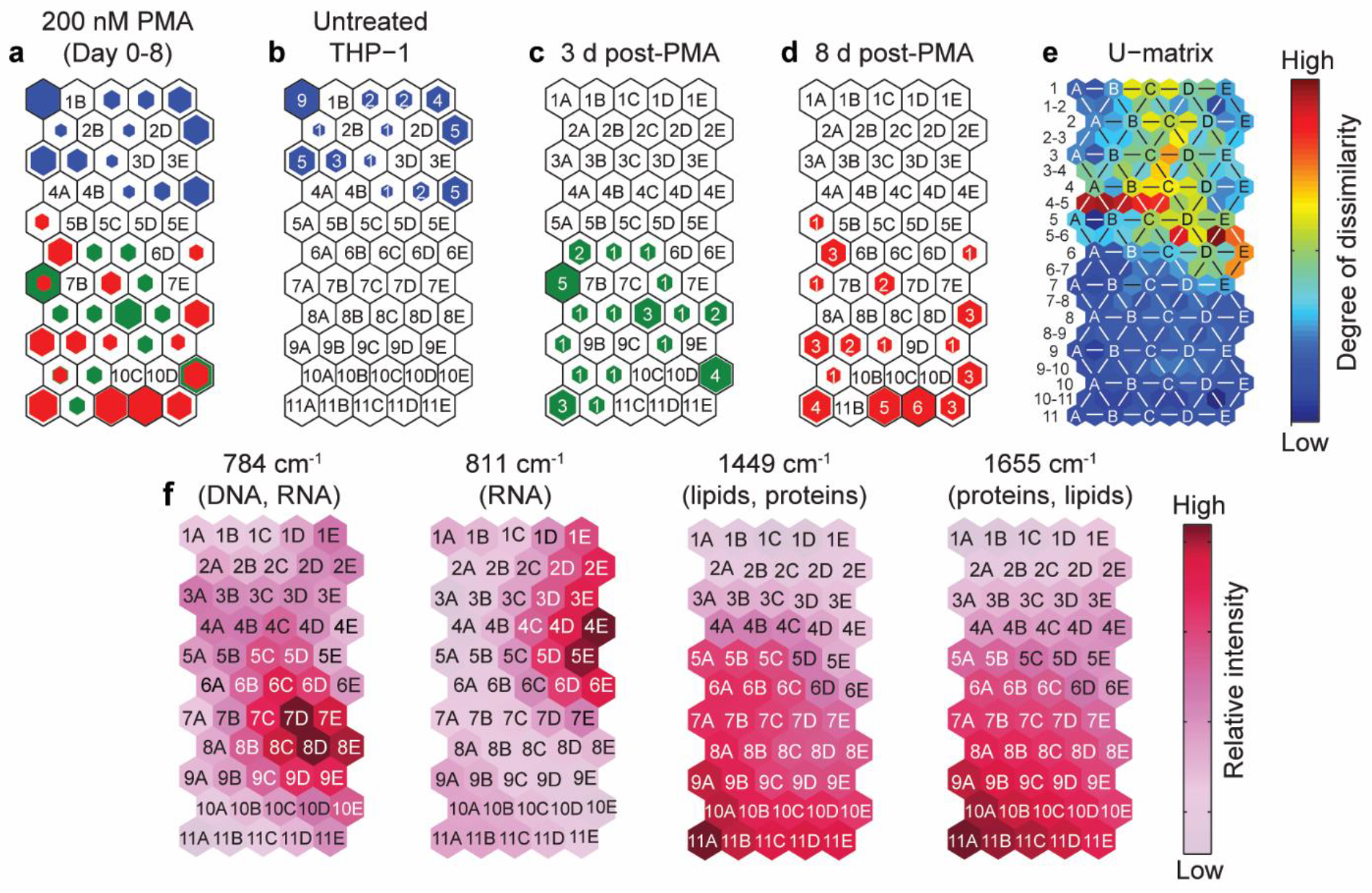
(a) Combined hit histogram shows the BMUs for THP-1 cells on poly-L-lysine-coated substrates at different times after treatment with 200 nM PMA. The size of the colored hexagons correlates with the number of spectra from untreated monocytes (blue), 3d post-PMA cells (green) and 8d post-PMA cells (red) assigned to the corresponding map units. (b-d) Hit histograms show BMUs for untreated THP-1 monocytes (b), 3d post-PMA cells (c), and 8d post-PMA cells (d). Colored hexagons are labeled with the number of hits. The number and letter on the empty hexagons denote their row and column, respectively. (e) Color-coded hexagons in the U-matrix encode the magnitude of spectral dissimilarity between individual neighboring map units (labeled with −, /, \), and the average dissimilarity between a map unit and all its neighbors (labeled with a letter). Letters refer to the column where the corresponding map unit is located in the hit histogram. Numbers located to the left of the plot denote the row where the corresponding map unit is positioned in the hit histogram. Hexagons labeled with lines (−, /, \) represent the dissimilarity between the map units in the hit histogram with the coordinates of the hexagons on either side of the line. (f) Component planes showing relative peak intensities corresponding to contributions from DNA/RNA (785 cm^−1^), DNA (1092 cm^−1^), lipids (primarily), proteins and other biomolecules (1449 cm^−1^), and proteins (primarily) and lipids (1655 cm^−1^).

During differentiation, the cell synthesizes the new biomolecules it will need to perform its new functions, and degrades the biomolecules that are no longer needed. These changes in biomolecular composition are expected to induce subtle differences between the Raman spectra of differentiated and undifferentiated THP-1 cells, which is reflected in the hit histograms in Figure 2a - d. The BMUs for the 41 spectra acquired from untreated THP-1 cells are confined to 13 adjacent map units located in rows 1 - 4 of the SOM, where rows 1 and 2 contain a higher percentage of BMUs for the untreated cell spectra than rows 3 and 4. In fact, row 1 contains more BMUs for the spectra of untreated THP-1 cells than any other row in the map. This suggest the map units in row 1 represent spectra that are most similar to the spectra of undifferentiated THP-1 cells. In contrast, the BMUs for the spectra acquired from THP-1 cells 8 d post-PMA treatment are 6 mostly adjacent map units at the other side of the map (rows 5 – 11) (Figure 2d). Notably, the last row of the SOM contains more BMUs for the spectra of THP-1 cells 8 d post-PMA treatment than any other row, which implies the map units in row 11 represent spectra that are most similar to those of completely differentiated THP-1 cells. Moreover, the relative positions of the BMUs for the untreated cell spectra and the 8 d post-PMA cell spectra suggests the extent of cell differentiation increases as the row of its BMU increases.

The BMUs for the Raman spectra of cells 3 days post-PMA treatment largely are adjacent to (23 of 30) or on the same map units (7 of 30) as the BMUs for the 8 d post-PMA spectra (Figure 2a – d). None of the BMUs for the 3 d post-PMA spectra are adjacent to or on the same map units as spectra from untreated THP-1 cells. This distribution indicates that the biochemistries of the 3 d post-PMA cells had changed enough to be spectrally distinct from the parent population, and more similar to cells subjected to PMA treatment and 5 days of rest. However, the BMUs for the 3 d post-PMA cell spectra are equally split between rows 5 – 7, 8 – 9, and 10 – 11, whereas the percentage of the BMUs for the 8 d post-PMA cell spectra in the same three regions increases down the SOM (20%, 25%, and 55% in rows 5 – 7, 8 – 9, and 10 – 11, respectively). The higher proportion of 8 d post-PMA spectra in higher row numbers indicates that these cells are spectrally, and therefore, biochemically, more different from untreated monocytes than are 3 d post-PMA cells. This is consistent with the previous finding that cells stimulated with PMA for 3 days have not finished differentiating from a monocyte-like phenotype to a macrophage-like phenotype.^8^

To gain insight into the extent of spectral variance between BMUs for THP-1 cells at different stages of PMA-induced maturation, we examined the U-matrix, which uses a color-coded scale to represent spectral variance (Figure 2e). The rows that correspond to the first four rows of the hit histogram, which contain the BMUs for the untreated THP-1 cells, exhibit higher variation than that corresponding to the rest of the hit histogram. The highest inter-unit spectral variation is located between the rows 4 and 5 of the hit histogram (row 4 – 5 in the U-matrix, Figure 2e), which contain the BMUs for untreated cells and a fully differentiate THP-1 macrophage, respectively. This signifies substantial differences between the spectral signatures, and thus biochemistries and phenotypes, of THP-1 cells before and after treatment with PMA. The hexagons in the U-matrix that correspond to BMUs for the PMA-treated cells in rows 7 through 11 of the hit histogram are mainly dark blue, indicating that these map units, and therefore the cells assigned to them, are spectrally very similar to one another. This supports our earlier conjecture that 3 d post-PMA cells exhibit many spectral hallmarks of the macrophage-like phenotype. Examination of the component planes (Figure 2f) reveals that the peaks at 1449 and 1655 cm^−1^, which are mainly produced by lipids and proteins,^24^ are higher in the BMUs for cells stimulated with PMA than in the BMUs for the untreated cells. This likely reflects the increase in lysosomes and mitochondria that is observed during differentiation.^8^ In comparison, a clear trend is not apparent in the component planes for the peaks (784 and 811 cm^−1^) associated with DNA and RNA (Figure 2f).

### Fibronectin does not affect THP-1 differentiation induced by high PMA concentration

Fibronectin modulates the maturation of primary monocytes into macrophages,^15^ so we explored the effect of fibronectin on PMA-triggered differentiation of THP-1 monocytes. Again, Raman spectra were acquired from THP-1 cells on fibronectin-coated substrates before PMA treatment, and at 3 or 8 days after the start of PMA treatment. The hit histograms (Figure 3a – d) show the BMUs for the spectra of untreated THP-1 cells and 8 d post-PMA cells were segregated within different regions of the SOM. rows 1 – 3 contain the BMUs for THP-1 cell spectra before PMA stimulation, and rows 4 - 9 contain the BMUs for the spectra of PMA-treated cells. However, the majority (~70%) of the 8 d post-PMA cells are concentrated in the bottom two rows of the hit histogram (Figure 3d). Based on the low topographical error for this SOM (Table S1), the spectra from untreated cells with BMUs in rows 1 – 3 are significantly different from the spectra of the PMA-treated cells with BMUs in -the bottom rows in the hit histogram. Thus, the 8 d post-PMA cell spectra assigned to the bottom rows of the hit histogram likely represent the fully differentiated macrophage-like phenotype (Figure 3d). The spectra for the 3 d post-PMA cells were distributed nearly equally between the BMUs corresponding to putatively fully differentiated cells (rows 8 and 9), and map units fairly proximal to the BMUs for both undifferentiated and fully differentiated cells (rows 4-6). This distribution implies that some cells had transformed fully into THP-1 macrophages (map units 8A, 8B, 8D, 8F, 9A-9F) after 3 d of PMA stimulation, while others were at an intermediate stage in their transition to the macrophage-like phenotype.

**Figure 3:**
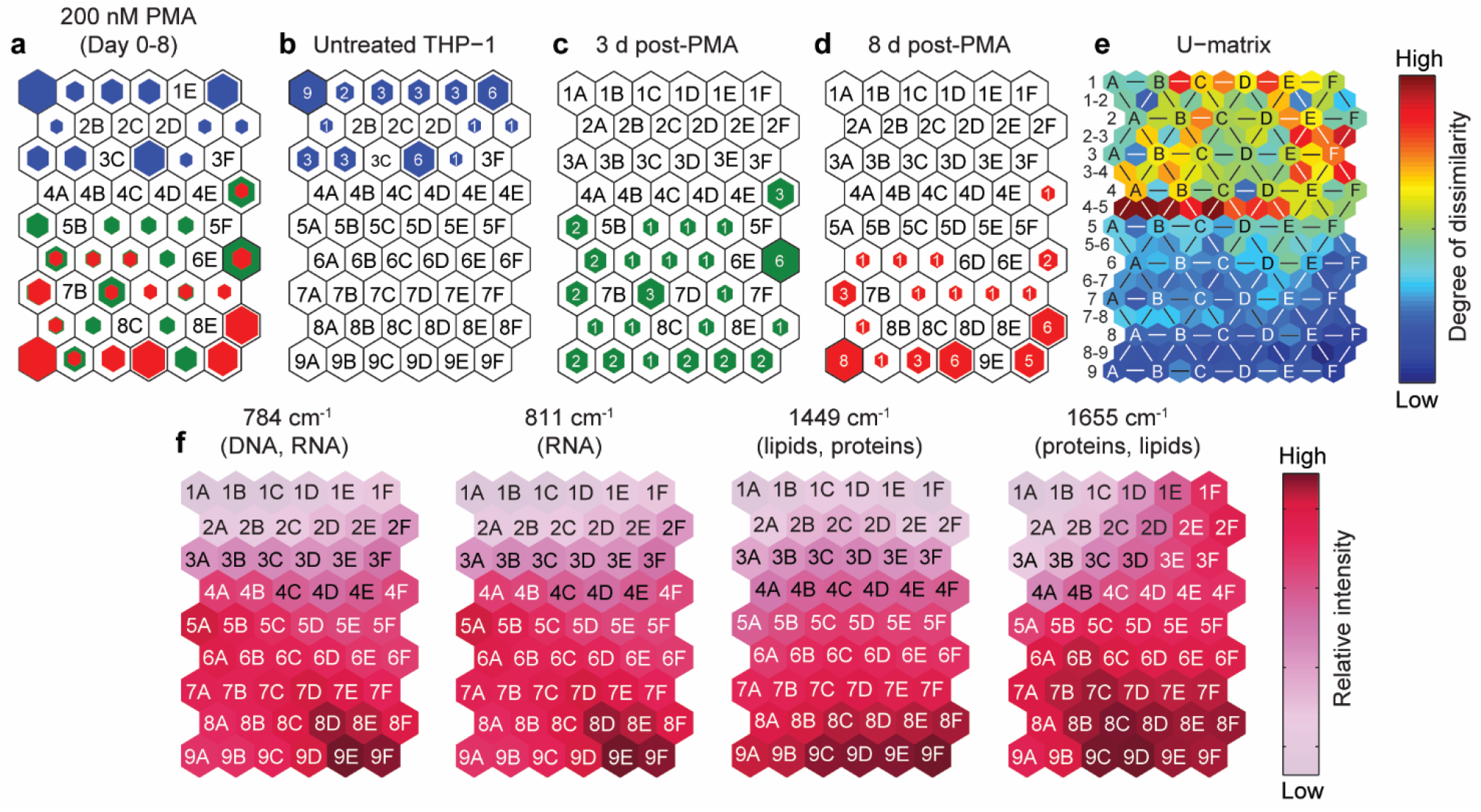
(a) SOM overlaid with hexagons showing number of hits for THP-1 cells on fibronectin-coated substrates before (blue) and after treatment with 200 nM PMA (green: 3d post-PMA; red: 8d post-PMA). (b-d) Hit histograms show BMUs for untreated THP-1 monocytes (b), 3d post-PMA cells (c), and 8d post-PMA cells (d). Numbers on the colored hexagons encode the number of spectra for which the underlying map unit was the BMU. Map units which were not BMUs for any spectra are labeled by their row number and column letter. (e) U-matrix uses color to show the degree of spectral dissimilarity between individual neighboring map units and the average dissimilarity between each map unit and all its neighbors. The markings on the U-matrix are explained in the Figure 2 caption. (f) Component planes showing relative peak intensities corresponding to contributions from DNA/RNA (785 cm^−1^), DNA (1092 cm^−1^), lipids (primarily), proteins and other biomolecules (1447 cm^−1^), and proteins (primarily) and lipids (1655 cm^−1^).

The levels of spectral variation between the three populations were consistent with the observations for cells on poly-L-lysine-coated substrates. The U-matrix (Figure 3e) shows orange and red hexagons that denote large spectral differences between the rows adjoining the BMUs for the untreated THP-1 cells in rows 1 - 3 and the post-PMA cells in rows 4 - 9. In addition, there is less spectral dissimilarity in the portion of the SOM that contains BMUs for PMA-treated cells than in the rows with untreated cells. Taken together, this suggests that the 3 d post-PMA cells are spectrally more similar to the 8 d post-PMA cells in rows 8 and 9 that presumably exhibit the macrophage-like phenotype than to undifferentiated THP-1 monocytes. The component planes showed trends in the peaks at 1655, 1449, 811, and 784 cm^−1^ that were similar to those observed for cells on poly-L-lysine stimulated with 200 nM PMA (Figure 3f). The similarity in these findings imply that the monocyte-macrophage transition induced in THP-1 cells on poly-L-lysine by high PMA concentration does not differ from that of cells on fibronectin.

### Fibronectin modifies the dynamics of THP-1 monocyte differentiation elicited by low PMA concentration

We hypothesized that stimulation with high PMA concentration might mask the effects of fibronectin on the dynamics of THP-1 cell differentiation to the macrophage-like phenotype. Because lowering the PMA concentration to ~10 ng/ml permitted the detection of macrophage response to weak subsequent stimuli,^11, 25^ we investigated whether fibronectin affects the extent of THP-1 monocyte differentiation induced by low PMA concentration as compared to the same stimulation performed in the presence of poly-L-lysine. Raman spectra were acquired from THP-1 cells in the presence of substrates coated with fibronectin or poly-L-lysine before treatment with 20 nM PMA, immediately after 3 d of PMA treatment, and after the PMA-treated cells were rested in PMA-free medium. The SOMs for the resulting spectra are shown in Figure 4. On poly-L-lysine substrates (Figure 4a – e), the BMUs for almost all the unstimulated and 8 d post-PMA cells are on opposite edges of the SOM (right and left, respectively, Figure 4a), indicating relatively significant spectral differences.

**Figure 4:**
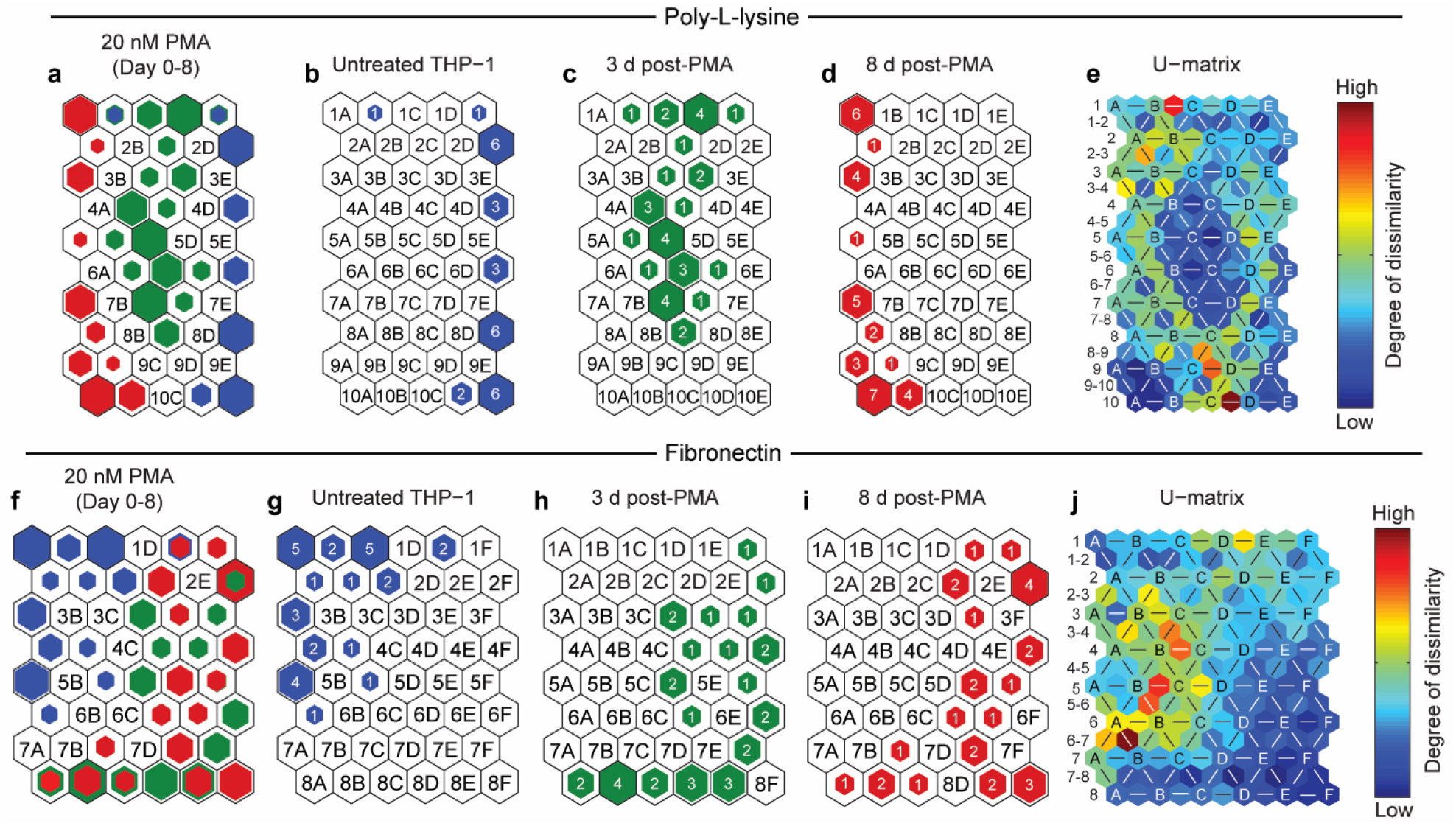
SOM shows hits for Raman spectra of THP-1 cells over 8 days after treatment with 20 nM PMA on poly-L-lysine (a-e) and fibronectin (f-j) coated substrates. Number of hits are encoded by the size of the colored hexagons overlaid on map units, with blue, green and red denoting untreated THP-1, 3 d post-PMA cells and 8 d post-PMA cell spectra respectively. (a, f) are combined hit histograms overlaid with hexagons showing the number of spectra from cells at all three time-points, (b, g) show BMUs for unstimulated THP-1 cells, (c, h) show hits corresponding to THP-1 cells treated with 20 nM PMA for 3 d, (d, i) show BMUs for THP-1 cells treated with 20 nM PMA for 3 d and allowed to rest for 5 d in PMA-free culture medium, (e, j) are the U-matrices showing the extent of dissimilarity between and within spectra assigned to individual map units as measured by the differences in peak intensities. The color of each hexagon is indicative of the amount of variance, as seen from the color bar next to the map. See Figure 2 caption for a detailed explanation of the markings on the map.

Notably, the BMU for one untreated cell spectrum (map unit 1B) is next to the BMUs for 8 d post-PMA cells (units 1A, 2A), but not other untreated cells. The magnitude of the spectral variance between map unit 1B and both 1A and 2A are denoted by blue hexagons in the U-matrix, signifying that the outlier untreated cell spectrum on map unit 1B is spectrally more similar to 8 d post-PMA cells than other untreated cells. This finding may indicate this cell spontaneously differentiated in the absence of PMA.

The hits for the spectra from 3 d post-PMA cells on poly-L-lysine-coated substrates are distributed between the BMUs for the untreated and 8 d post-PMA cells (columns B – D, Figure 4a - c). This implies stimulation with 20 nM PMA for 3 days produces cells with an intermediate stage of differentiation. None of the BMUs for the 3 d post-PMA cells on poly-L-lysine-coated substrates were on the same map units as the BMUs for untreated cells or 8 d post-PMA cells, indicating all the cells had undergone some differentiation, but none were fully differentiated. We also attempted to investigate whether THP-1 cells on poly-L-lysine reached the same extent of differentiation 8 d after PMA treatment regardless of whether 20 or 200 nM PMA was employed for stimulation by creating a single SOM of the spectra of THP-1 cells on poly-L-lysine (Figure S1). However, because the experiments with the low and high PMA concentrations were performed months apart, we were unable to separate the spectral dissimilarity indicative of differences in experimental conditions from those related to cell differentiation stage (see SI for details).

Unlike our observations for cells on poly-L-lysine BMUs for spectra from 3 d post-PMA cells on fibronectin-coated substrates overlapped substantially with the BMUs for spectra from 8 d post-PMA cells on fibronectin, irrespective of PMA concentration (Figure 4f - j). Approximately 40% of untreated THP-1 cell spectra were assigned to the top left region of the SOM (columns A - C of rows 1 and 2) and close to 30% were assigned to map units in column A of rows 3 through 5 (Figure 4g). The BMUs for the spectra from most 8 d post-PMA were on the right side of the SOM, though some BMUs span the entire bottom two rows (Figure 4i). Most of the spectra from the unstimulated and 8 d PMA-treated cells were assigned to different regions of the SOM, though some are positioned next to each other (map units 2C and 2D, 1E and 1F) or on the same units (map unit 1E). More than half (~56%) of the spectra collected from cells after 3 d have the same BMUs as 8 d PMA-treated spectra (map units 1F, 2F, 3E, 4F, 5F, 6D, 8A, 8B, 8C, and 8E). The rest of the BMUs for the 3 d post-PMA cell spectra are assigned to map units adjacent to the BMUs for 8 d post-PMA cell spectra. Moreover, fairly high spectral dissimilarity between untreated and PMA-treated cells is indicated by the green or yellow colored hexagons in the U-matrix that denote the spectral difference between the right-hand border of the clustered BMUs for the untreated cells and the neighboring map units to the right that are BMUs for post-PMA cells (Figure 4j). The U-matrix also shows 3 d post-PMA cells have low spectral dissimilarity among themselves and compared to the BMUs for 8 d post-PMA cells, but moderate to relatively high spectral dissimilarity compared to the BMUs for untreated cells (Figure 4j). Taken together, these results suggest that 3 d post-PMA cells are spectrally more similar to 8 d post-PMA cells than untreated cells. We also infer the THP-1 cells on fibronectin-coated substrates had not only begun differentiating after 3 d simulation with the lower PMA concentration, but some had advanced to a stage where their spectral features, and thus phenotypes, were very similar to those of the 8 d post-PMA cells.

Finally, we investigated whether the PMA concentration used for stimulation affects the extent of differentiation reached by THP-1 cells on fibronectin, by creating a SOM with spectra acquired from all the cells with fibronectin-coated substrates (Figure S1). The BMUs for the spectra acquired from 3 d and 8 d post-PMA-treated cells were on the same or neighboring map units, regardless of the PMA concentration used for stimulation (Figure S1). This indicates that in the presence of fibronectin, THP-1 cells reach approximately the same extent of differentiation following 3 d of simulation with either PMA concentration (see SI for details).

Overall, these results show that the PMA concentration used for stimulation affects the spectral features that encode for the extent of THP-1 cell differentiation when simulation is performed in the presence of substrates coated with poly-L-lysine. However, cells on fibronectin-coated substrates stimulated with low and high PMA concentration for 3 d had high spectral similarity, and thus had reached the same extent of differentiation. This implies that the substrate coating may affect the PMA-induced differentiation trajectory at lower levels of PMA stimulation.

## DISCUSSION

Macrophages are important immune cells, capable of fulfilling a range of functions from guiding the immune response and mediating tissue repair during injury to maintaining physiological homeostasis.^1, 26^ As a monocytic cell line that can be readily differentiated into macrophage-like cells with increased adherence, THP-1 cells have been routinely used as in vitro models to study macrophage polarization and how it contributes to the pathogenesis of diseases such as atherosclerosis^4^ and fibrosis. Differentiation into macrophages triggered by PMA produces macrophage-like cells which differ markedly in their phenotypic and metabolic attributes, based on the concentration of stimuli used^8, 25^, the length of incubation and whether or not cells were rested in culture media after withdrawal of stimuli.^8^

There is no consensus on the optimal level of PMA stimulation required for differentiation of THP-1 monocytes into macrophages or any of the subsequent polarized phenotypes.^8, 11, 25^ High concentrations are reported to produce resting macrophages that mirror the physiology and metabolism of macrophages derived from primary monocytes.^8^ In conflicting reports, stimulation with high PMA concentration induced a more mature macrophage phenotype^11, 25^ based on the over-expression of cytokines associated with polarized macrophages.^25^ In this study, we explored the effect of two different PMA concentrations on the differentiated macrophage-like phenotype based on the Raman spectral fingerprints of THP-1 cells. SOMs constructed from the spectra acquired from cells before and after treatment to map spectral changes related to transformation into macrophage-like cells. Our results show that the Raman spectra from cells stimulated with both low and high PMA concentrations had largely diverged from the untreated cells 3 days after treatment. At high PMA concentration (200 nM), the 3 d post-PMA cells tended to cluster with 8 d post-PMA cells regardless of the substrate coating, which suggests that many, though not all, of these cells had reached the THP-1 macrophage phenotype. Similarly, the cells on fibronectin-coated substrates stimulated with 20 nM PMA for 3 d also had substantial overlap with the 8 d post-PMA phenotype, indicating low PMA concentration combined with culture on fibronectin induces many of the biochemical changes necessary for transformation into THP-1 macrophages. However, cells on poly-L-lysine showed a more graded response to 20 nM PMA, and less overlap between the BMUs for spectra taken 3 and 8 days after treatment. This implies that treatment with 20 nM PMA on poly-L-lysine produces cells of an intermediate nature, which have distinct spectral characteristics from the macrophage-like cells purported to result after incubation with 200 nM.

Of note, adherence is generally accepted as a measure of the extent of THP-1 differentiation into macrophages. Consequently, treated cells that do not wash off after treatment are assumed to have differentiated, often without verification of their state with immunofluorescence. In this study, the spectra from PMA-treated cells were taken from adherent cells, yet the SOM showed some spectral and thus biochemical differences between 3 d post-PMA and 8 d post-PMA cells, especially for cells on poly-L-lysine treated with 20 nM PMA. This suggests that attachment is not a good indicator of the fully differentiated macrophage-like phenotype. The phenotypes of adherent THP-1 cells should be validated with a complementary technique, which would help ensure that results obtained from different THP-1 cell experiments are comparable and reproducible.

The cell-adhesive motifs present in ECM proteins have been implicated in monocyte-macrophage transformation and subsequent polarization in vitro and in vivo.^15, 27–29^ Furthermore, increased uptake of proteins by endocytosis, and elevated expression of macrophage-associated markers and intracellular enzymes were observed in PBMCs after 5 d of culture on slides coated with fibronectin and collagen I as compared to those on collagen IV.^15^ This finding that fibronectin modulates primary monocyte differentiation is consistent with our conclusion that THP-1 cells on fibronectin that had been stimulated with 20 nM PMA are more differentiated than THP-1 cell subjected to the same PMA simulation conditions in the presence of substrates coated with poly-L-lysine. The closer proximity of the BMUs of cell spectra acquired 3 d and 8 d post-20 nM PMA treatment observed for substrate coating with fibronectin but not poly-L-lysine suggests that fibronectin hastened the differentiation of THP-1 cells in response to 20 nM PMA. High PMA concentrations seemed to mask this effect, as seen in the similar proximities of the BMUs for cells 3 d and 8 d post-treatment with 200 nM PMA, regardless of substrate coating. Interestingly, high PMA concentration (> 100 ng/ml) was reported to differentiate THP-1 monocytes past the resting macrophage state based on their insensitivity to polarizing stimuli.^11^ Our observation that high PMA concentrations mask the effects of fibronectin on THP-1 maturation agrees with the prior finding that higher PMA concentrations can mask the effects of other external factors.

## CONCLUSIONS

Based on self-organizing maps of single-cell Raman spectra acquired from living cells after exposure to different stimuli, we discovered fibronectin increases the extent of THP-1 monocyte differentiation into macrophage-like cells reached at a specified time following stimulation when low, but not high concentrations of differentiation-inducing stimuli are used. Future work that follows the expression of genes and markers elicited by culture on ECM-coated substrates may delineate the mechanistic details underlying the change in behavior of THP-1 cells on fibronectin.

## ACKNOWLEDGEMENTS

We thank Isamar Pastrana-Otero for fabricating the gold-coated substrates used for Raman spectroscopy in this work. We are also grateful to Manjari Mishra and Dr. Shobhna Kapoor at the Indian Institute of Technology Bombay, India for helpful discussions on culturing and differentiating THP-1 cells. Raman microspectroscopy was performed in the Microscopy Suite of the Imaging Technology Group at the Beckman Institute for Advanced Science and Technology, University of Illinois. The steps involved in substrate fabrication were performed in the Frederick Seitz Materials Research Laboratory, University of Illinois. This work was partially supported by the National Institute of Heart, Lung and Blood Institute under grant number R21 HL132642. The authors also acknowledge additional funding provided by the Dept. of Chemical and Biomolecular Engineering at the University of Illinois.

