## Supplemental Material for "Exploring the maturation of a monocytic cell line using self-organizing maps of single-cell Raman spectra"

### Supplemental Table 1: SOM quality measures for all the maps generated

| PMA concentration | Substrate coating | mqe | te |
|---|---|---|---|
| **200 nM** | Combined | 21.015 | 0.004 |
| | Poly-L-lysine | 20.195 | 0 |
| | Fibronectin | 21.693 | 0 |
| **20 nM** | Combined | 19.378 | 0.016 |
| | Poly-L-lysine | 19.053 | 0.011 |
| | Fibronectin | 19.427 | 0.011 |
| **20 and 200 nM** | Poly-L-lysine | 20.234 | 0.010 |
| **20 and 200 nM** | Fibronectin | 21.100 | 0 |
| **20 and 200 nM (combined)** | Combined | 20.883 | 0.007 |

Table S1: Summary of map statistics for different combinations of PMA concentration and substrate chemistry

**Supplemental Table 2: Quantifying the spread of BMUs for each cell population under all conditions studied in this work**

| PMA concentration (nM) | Substrate coating | Treatment group | Mean inter-BMU distance (within-group spread) |
|---|---|---|---|
| **200 nM** | Combined | Untreated THP-1 | 3.478 |
| | | 3 d post-PMA | 3.868 |
| | | 8 d post-PMA | 3.567 |
| | Poly-L-lysine | Untreated THP-1 | 1.956 |
| | | 3 d post-PMA | 3.846 |
| | | 8 d post-PMA | 3.311 |
| | Fibronectin | Untreated THP-1 | 2.242 |
| | | 3 d post-PMA | 3.691 |
| | | 8 d post-PMA | 3.759 |
| **20 nM** | Combined | Untreated THP-1 | 5.060 |
| | | 3 d post-PMA | 3.114 |
| | | 8 d post-PMA | 4.716 |
| | Poly-L-lysine | Untreated THP-1 | 3.027 |
| | | 3 d post-PMA | 3.116 |
| | | 8 d post-PMA | 4.460 |
| | Fibronectin | Untreated THP-1 | 2.190 |
| | | 3 d post-PMA | 3.152 |
| | | 8 d post-PMA | 3.535 |
| **20 and 200 nM (combined)** | Combined | Untreated THP-1 | 4.170 |
| | | 3 d post-PMA | 4.230 |
| | | 8 d post-PMA | 5.261 |
| | Poly-L-lysine | Untreated THP-1 | 1.360 |
| | | 3 d post-PMA | 3.332 |
| | | 8 d post-PMA | 3.023 |
| | Fibronectin | Untreated THP-1 | 2.130 |
| | | 3 d post-PMA | 3.623 |
| | | 8 d post-PMA | 3.621 |

Table S2: Average within-population spread quantified by the distance between BMUs for cells at each time-point.

**Supplemental experiment 1: Comparison of cells on substrates with the same coating that were stimulated with high and low PMA**

We investigated whether the 8 d post-PMA cells treated with 20 nM PMA were less differentiated than cells treated with 200 nM PMA. SOMs were constructed of spectra from THP-1 cells treated with both PMA concentrations on the same coatings (poly-L-lysine, Figure S1a – e; fibronectin, Figure S1f - j). The relative distributions of the cell spectra on the SOM differed between the two substrate coatings. Surprisingly, the untreated cell spectra on poly-L-lysine form two distinct clusters in the central and top-left regions of the map (light blue and dark blue, respectively, in Figure S1b). Because these samples consisted of untreated cells on the same coating, but they were prepared several months apart, this spectral variation likely reflects differences in experimental conditions (i.e., number of passages or length of culture on substrates before spectral acquisition). The spectra from the 3 d post-PMA cells subjected to both PMA levels were assigned to overlapping or neighboring units in the bottom three rows of the map, although the BMUs for a few cells extend up to row 4 (column G, Figure S1c). This indicates the 3 d post-PMA cells had similar spectral features regardless of the PMA concentration used for stimulation. Notably, the spectra of a few cells 3 d post treatment with 20 nM PMA were assigned to map units that were the BMUs for untreated cells (map units 7A, 9A, Figure S1c), indicating little differentiation had occurred under these conditions.

Conversely, 8 d post-PMA cells stimulated with different PMA concentrations were mapped to non-overlapping regions of the SOM (Figure S1d). Most spectra taken from cells 8 d after treatment with 200 nM PMA were assigned to contiguous map units in the bottom row (red hexagons, Figure S1d). Similar to the SOM shown in Figure 2, several of these map units were also the BMUs for spectra of cells acquired after 3 d of PMA stimulation (Figure S1a, c). Again, this implies that a fraction of cells treated for 3 d with 200 nM PMA had completely transformed into THP-1 macrophages. Interestingly, spectra from 8 d post-PMA cells treated with 20 nM PMA were split into two groups of BMUs on the top-right and bottom-left edges of the SOM (orange hexagons, Figure S1d). The U-matrix indicates that the cluster of BMUs on the top-right is spectrally dissimilar to the nearest BMUs for other cell populations (Figure S1e). Because these cells don’t overlap with the BMUs for 8 d post-200 nM PMA cells, nor do they lie between untreated and 8 d post-PMA cells, it is unlikely that these cells are at an intermediate stage in the differentiation trajectory. This, along with the lack of overlap with cells 8 d post treatment with 200 nM PMA, indicates that these two cell populations are spectrally distinct, which may signify differences in cell biochemistry induced by the PMA concentration used for stimulation.

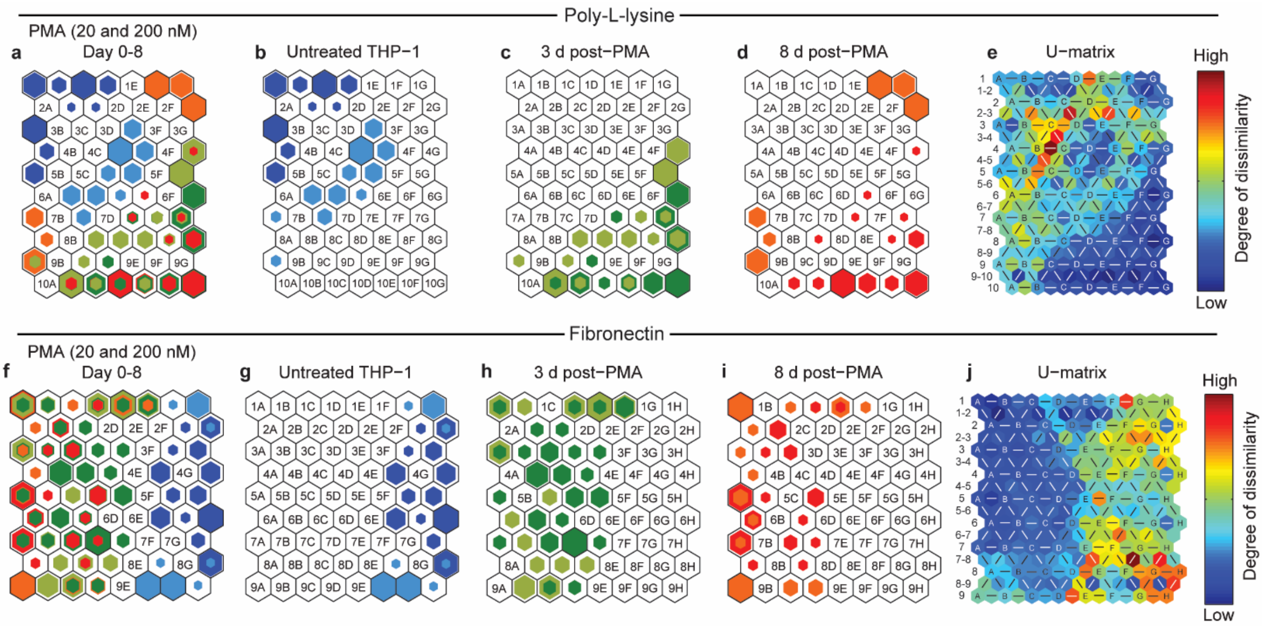


Figure S1: (a) Hit histograms overlaid with BMUs for THP-1 cells cultured on poly-L-lysine (a-e) and fibronectin-coated (f-j) substrates at different time-points after treatment with 200 nM PMA (lighter colors) and 20 nM PMA (darker colors). The size of the colored hexagons is correlated with the number of spectra from untreated monocytes (blue), 3d post-PMA cells (green) and 8d post-PMA cells (red) assigned to the corresponding map units. (b-d) Individual hit histogram showing BMUs for untreated THP-1 monocytes (b,g), 3 d post-PMA cells (c,h), and 8 d post-PMA cells (d,i). (e,j) U-matrix reveals the amount of spectral variation between neighboring map units using a color scale. See Figure 2 caption for a detailed explanation of the markings on the map.

For stimulation performed in the presence of fibronectin-coated substrates, the differences between spectra acquired from cells treated with the two PMA levels are less significant (Figure S1f-i). The BMUs for the untreated cells on substrates coated with fibronectin were confined to three columns on the right side of the map (Figure S1g). In contrast, the distribution of 8 d post-PMA cell spectra was skewed to the left, although individual spectra were assigned to units spread over a larger region of the map (Figure S1i). Similarly, cells treated for 3 d with PMA (Figure S1h) were assigned to the central-left region and shared half of their BMUs (18 out of 34) with 8 d post-PMA cells. Importantly, the spread was not dictated by the level of PMA stimulation, as cells treated with 20 and 200 nM PMA were often assigned to the same or neighboring map units, which indicates that the spectral changes accompanying the differentiation process have converged after 3 d of PMA treatment for cells cultured on fibronectin-coated substrates. A few of these BMUs were on a map unit immediately adjacent to the BMU for an unstimulated cell spectrum (map unit 1F, Figure S1h). Although they are located next to each other on the SOM, the U-matrix shows high spectral dissimilarity between these map units.

### Supplemental experiment 2: Combined SOM of spectra from THP-1 cells on both substrate coatings stimulated with both PMA concentrations

An SOM was constructed using the spectra from cells subjected to the two concentrations of PMA. BMUs for unstimulated and putative differentiated (8 d post-PMA) cells were separated by the diagonal boundary formed by map units 10A, 9C, 8E, 7F and 6G. As expected, the unstimulated cells were assigned to similar or adjacent map units in the upper left of the SOM. The treated cells showed relatively less overlap with most of the cells subject to the higher concentration of PMA assigned to map units farther right, but the extent of overlap varied with the strength of PMA stimulus. Cells stimulated with the 8-day protocol at lower PMA levels were more likely to be assigned on or close to the diagonal boundary. A few of the BMUs were located in a region close to hits for undifferentiated cells (map units 1E, 2E, 3E). These results suggest that stimulation with 20 nM PMA is not sufficient to reliably induce the required spectral and phenotypic changes characteristic of macrophages in all of the treated cells. The few PMA-treated cells present on or close to the diagonal boundary (map units 10A, 11A, 12A and 7C) can be reasoned to have a more intermediate phenotype compared to cells assigned to the right (column H, rows 8-13) and bottom (map units 13D-H) of the SOM.

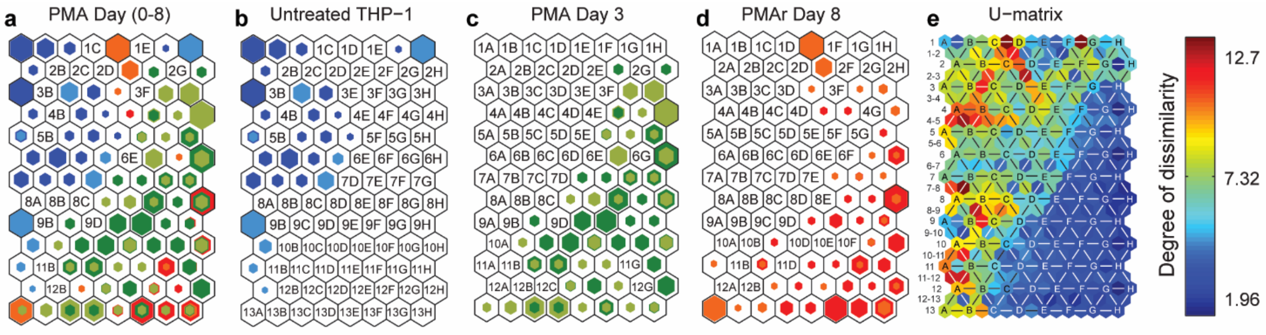


Figure S2: Hit histograms combining Raman spectra obtained from cells subjected to 20 nM (lighter colors) and 200 nM PMA (darker colors)-coated substrates showing BMUs for a) combined hit histogram overlaid with BMUs for all three treatments, b) unstimulated THP-1 cells, c) THP-1 cells treated with 20 nM (light green) or 200 nM (dark green) PMA for 3 days, d) THP-1 cells treated with 20 nM (orange) or 200 nM (red) PMA for 3 days and allowed to rest for 5 days in PMA-free culture medium. e) The corresponding U-matrix. See the caption for Figure 2 (main text) for a detailed explanation of the markings on the map.
